# Reduced feedback barely slows down proprioceptive recalibration

**DOI:** 10.1101/2022.01.19.476977

**Authors:** Jennifer E. Ruttle, Bernard Marius ’t Hart, Denise Y. P. Henriques

**Affiliations:** Centre for Vision Research, York University, Toronto, Canada; Department of Psychology, York University, Toronto, Canada; School of Kinesiology and Health Science, York University, Toronto, Canada

## Abstract

Introducing altered visual feedback of the hand results in quick adaptation of reaching movements. And while this may be partly due to explicit strategies, our lab has shown that implicit changes like reach aftereffects and shift in estimates of the unseen hand, can also emerge and even saturate within a few training trials. The goal of the current study is to determine whether these rapid changes in unseen hand position that occur during classical visuomotor adaptation are diminished or slowed when feedback during training is reduced. We reduced feedback by either providing visual feedback only at the end of the reach (terminal feedback) or constraining hand movements to reduce efferent contribution (exposure). We measured changes as participants completed reaches with a 30° rotation, a −30° rotation and clamped visual feedback, with these two “impoverished” training conditions, along with classical visuomotor adaptation training, while continuously estimating their felt hand position. Classic continuous-cursor training produced exemplary learning curves and rapid and robust shifts in felt hand position. Training with terminal feedback slightly reduced the initial rate of change in overall adaptation and but not the magnitude of shifts in felt hand position. Finally using a robot to constrain and deviate hand movement direction, called exposure training, only delayed saturation of proprioceptive changes by a single trial and these changes were slightly smaller than those during classical training. Taken together, adaptation and shifts in felt hand position are a rapid and robust responses to sensory mismatches and are only slightly modulated when feedback is reduced. This means that, given a visuo-proprioceptive mismatch, the resulting shift in sense of limb position can contribute to movements from the start of adaptation.

## Introduction

Our lab has recently demonstrated that changes in direct measures of implicit learning, such as reach aftereffects and shifts in estimates of the unseen hand position, during visuomotor adaptation develop surprisingly quickly. We find reach aftereffects and changes in estimates of hand location saturating within 3 and 1 training trials respectively. In this study, we aim to better understand and characterize this change in hand localization during adaptation but measuring whether the speed by which these localization shifts saturate during adaptation can be reduced when feedback during adaptation is also diminished. To do so, we use two common paradigms with reduced visual feedback that still leads to visuomotor adaptation: terminal feedback and cross-sensory exposure. We characterize both the rate of adaptation and shift in hand localization in these two paradigms and compare it with that produced during classical visuomotor adaptation.

In terminal feedback training the cursor representing the unseen hand is provided only at the end of the reach movement. Reducing visual feedback to the end of the reach during visuomotor rotation training has been shown in some studies to reduce the extent of learning and the magnitude of reach aftereffects (Barkley, Salomonczyk, Cressman, & Henriques, 2014; Hinder, Riek, Tresilian, De Rugy, & Carson, 2010; Hinder, Tresilian, Riek, & Carson, 2008; Taylor, Krakauer, & Ivry, 2014) although this is not always the case (Brudner, Kethidi, Graeupner, Ivry, & Taylor, 2016; Heuer & Hegele, 2008; Rand & Rentsch, 2016). Whether terminal feedback also affects the rate of adaptation is usually not quantified. Compared to continuous cursor feedback, training with terminal feedback has been shown to also reduce or slow down the changes in estimate of hand location (Barkley et al., 2014; Izawa & Shadmehr, 2011) but the rate of change has not been determined on a trial-by-trial basis.

Cross-sensory exposure training involves either passively moving the unseen hand or using a force-channel that deviates its direction, while the cursor moves directly to a target. Despite minimizing the motor or efferent signals involved, this passive exposure to a discrepancy between seen and felt hand location leads to similar or smaller but significant reach aftereffects (Cressman & Henriques, 2010; Mostafa, ’t Hart, & Henriques, 2019; Ruttle, ’t Hart, & Henriques, 2018; Salomonczyk, Cressman, & Henriques, 2013) and can facilitate subsequent adaptation to the same perturbation in a classic visuomotor paradigm (Bao, Lei, & Wang, 2017; Sakamoto & Kondo, 2015; Tays, Bao, Javidialsaadi, & Wang, 2020). Not surprisingly, such training also leads to changes in hand localization, which are similar in size to those elicited when the reaches are self-generated during classical visuomotor adaptation. This suggests that this proprioceptive recalibration is primarily driven by the visual-proprioceptive mismatch between the hand and the cursor. We have previously measured hand localization shifts on a trial-by-trial basis, allowing us to assess the rate of change for these shifts. Given these shifts in hand localization saturates within a single trial during classical visuomotor adaptation, our aim was to determine if a similar saturation rate occurs when the motor system is less engaged.

While it is reasonable to assume that reducing and removing availability of certain types of feedback, like that for terminal feedback or exposure training, should affect the time-course and/or asymptotic level of adaptation, this is far from settled. Moreover, it is unknown whether reducing this feedback can also slow down the rapid saturation of shifts in hand localization. Our goal is to qualify and model the rate by which these changes in felt hand position saturate on a trial-by-trial basis and how they compare across exposure, continuous or terminal feedback training. By measuring shifts in felt hand position after every training trial with these three feedback types, we can identify the role feedback has during ongoing adaptation and proprioceptive recalibration.

## Methods

### Participants

96 (mean age=22.17, range=18-46, males=22) right-handed, healthy adults participated in this study, and gave prior, written, informed consent. All procedures were in accordance with institutional and international guidelines and were approved by the York Human Participants Review Subcommittee.

### Apparatus

The experimental set-up is illustrated in Fig 1A. While seated, participants held a vertical handle on a two-joint robot manipulandum (Interactive Motion Technologies Inc., Cambridge, MA, USA) with their right hand such that their thumb rested on top of the handle. A reflective screen was mounted horizontally, 14 cm above the robotic arm. A monitor (Samsung 510 N, 60 Hz) 28 cm above the robotic arm presented visual stimuli via the reflective screen to appear in the same horizontal plane as the robotic arm. A Keytec touchscreen 2 cm above the robotic arm recorded reach endpoints of the left hand, to unseen, right hand targets (see (Cressman & Henriques, 2009) for more details). Subject’s view of their training (right) arm was blocked by the reflective surface and a black cloth, draped between the touch screen and their right shoulder. The untrained, left hand was illuminated, so that any errors in reaching to the unseen, right target hand could not be attributed to errors in localizing the left, reaching hand.

**Figure 1.**
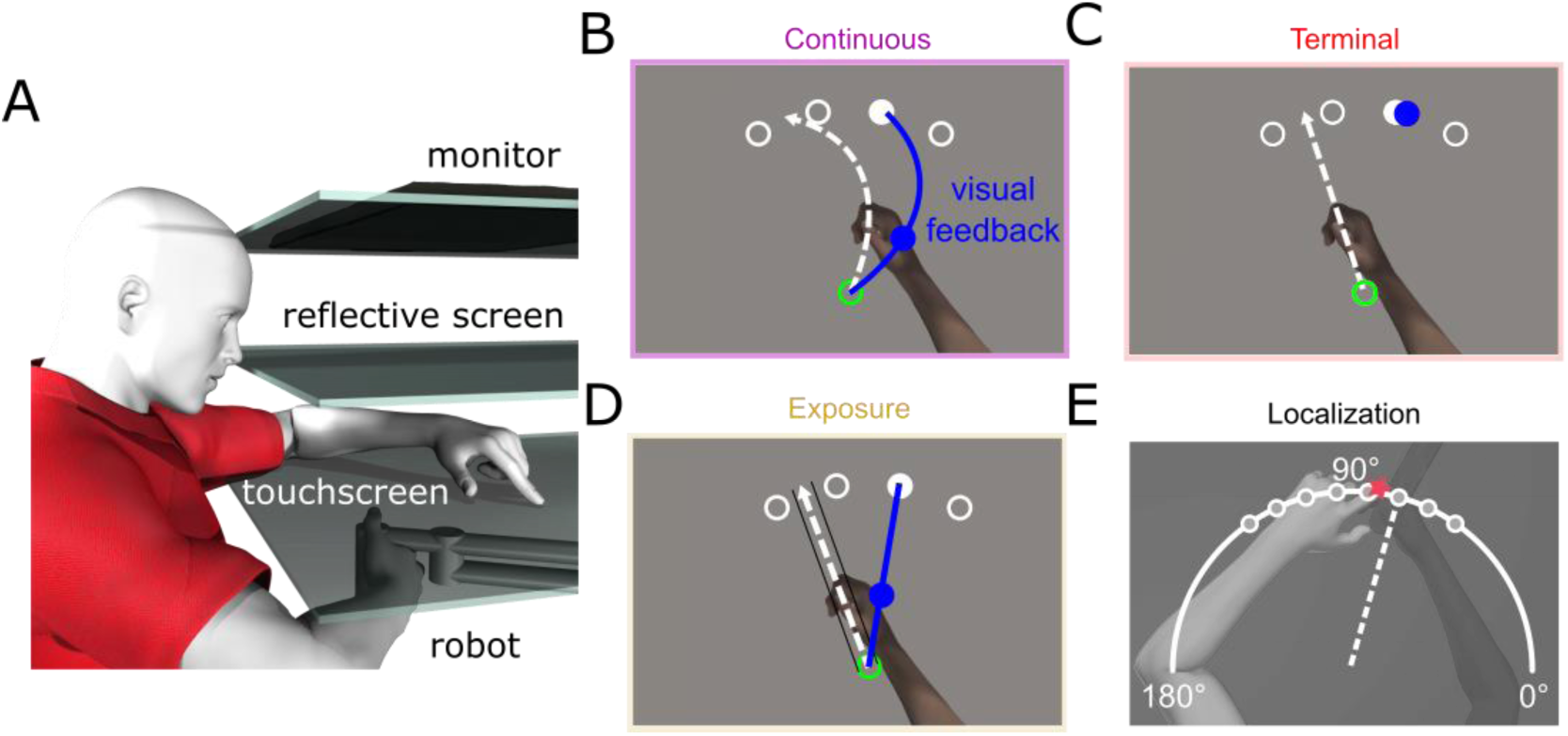
Experimental setup and design. **A:** Side view of the experimental set-up. The top layer is the monitor, middle layer is the reflective screen, and the bottom opaque layer is the touchscreen. The robot is depicted beneath with the participants’ right hand grasping it. B-D: Top views of task specific set-ups. **B:** Continuous training trial. The home position is represented by a green circle with a 1 cm diameter; located approximately 20 cm in front of the subject. Targets are represented by white circles with a 1 cm diameter located 12 cm radially from the home position at 60°, 80°, 100° and 120°. Participants hand cursor was also a 1 cm diameter blue circle. **C:** Terminal training trial. The same hand cursor was only visible at the end of the movement for 500 ms to allow for comparison to the visible target. **D:** Exposure training trial. The robot constrained the participants movements (denoted by solid black lines either side of white dashed line), so they perfectly countered the rotation and only decided the distance they moved. **E:** Localization test trial. Participants were passively moved to one of the eight target locations, 55°, 65°, 75°, 85°, 95°, 105°, 115° and 125°. Subsequently, participants used a touch screen to indicate on a white arc spanning 180° where their unseen right hand was.

### Stimuli

At the beginning of each trial, we displayed one of four potential targets, white 1 cm diameter circles, 12 cm from the start position at 60°, 80°, 100° and 120°. The home position (green 1 cm circle) and the participants hand cursor (blue 1 cm circle) were also visible at the beginning of the trial (for 2 of the 3 paradigms). During proprioceptive localization trials a white arc, 12 cm from the home position, was visible on the screen spanning from 0° to 180°. Participants were required to hold their hand still at the home position for 250 ms before any trial would begin.

### Trial Types

#### Classic training trials

Participants (N=32) reached as accurately as possible with their right hand to one of four possible target locations, while their hand cursor was continuously visible (Fig 1B). In all reaching trials, i.e., with cursor and with clamped cursor, participants had to reach out 12 cm from the home position to a force cushion within 800 ms. Participants received auditory feedback throughout training indicating if they met the distance-time criteria or not. The target would then disappear, and the robot manipulandum returned the right hand to the home position where they waited 250 ms for the next trial. The hand cursor was aligned with the hand for the first 64 training trials, then rotated 30° CW for 160 training trials and then rotated 30° CCW for 16 training trials. This was followed by 48 error-clamped trials, dashed lines in Fig 2, which were identical to the reach training trials except that the cursor always moved on a straight line to the target. The distance of the error-clamped cursor from the home position was identical to the distance of the hand from the home position.

**Figure 2.**
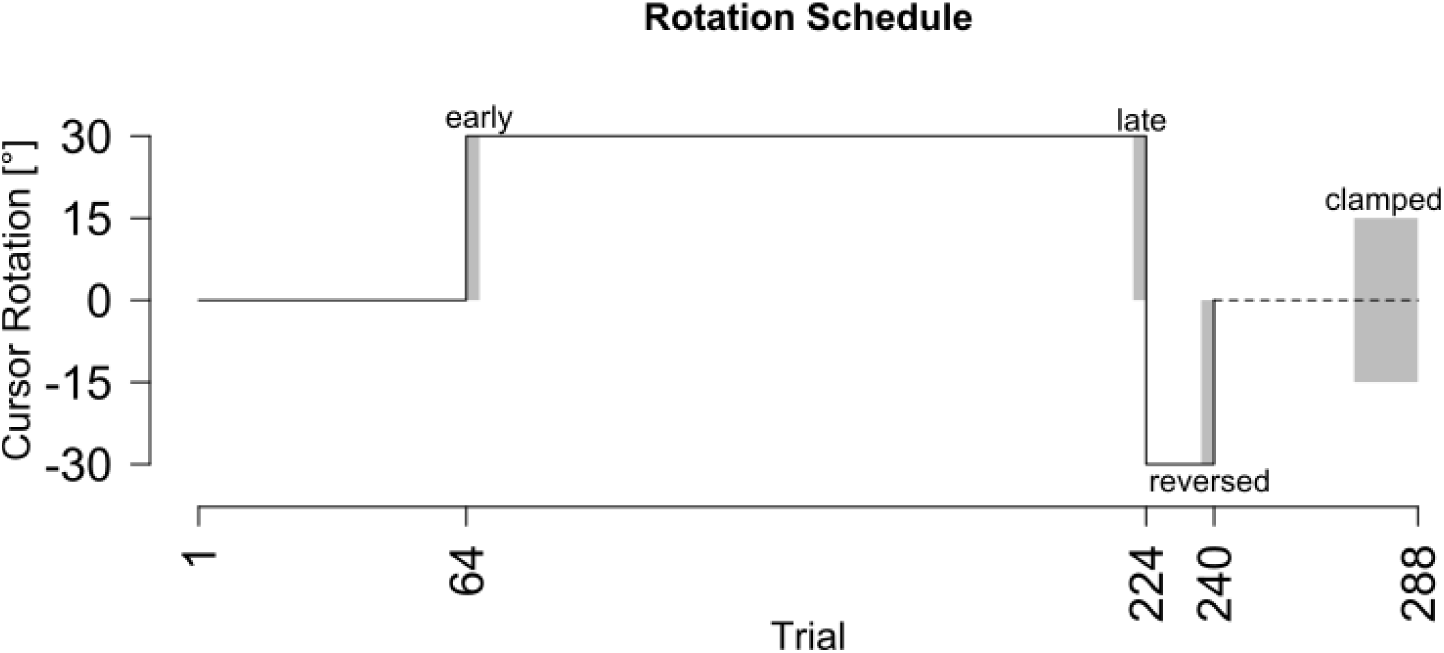
Experimental Schedule. Participants reached to visual targets with a perturbation denoted by the black line. The dotted line at the end of the paradigm signifies clamped trials where there was no visual error as the cursor always moved to the target regardless of the participants movement direction. Trials included in analysis are as follows: early=trials 65-68; late=trials 221-224; reversed=trials 237-240; clamped=273–288.

#### Terminal training trials

Terminal training trials were identical to classic training trials, except that the participants’ (N=32) hand cursor was not visible during the entire reach movement, from the home position to the target (Fig 1C). Once the participant moved their hand 12 cm from the home position, the robot locked their hand in place and the hand cursor became visible for 500 ms for the participant to be able to see any potential movement errors. The auditory cues were present to encourage consistent speed throughout the experiment. These participants also experienced a phase of error clamped trials which were identical to the classical clamp trials, with the cursor being visible the entire trial, not just at the end.

#### Exposure training trials

Exposure training trials differ from those in the previous two paradigms (Fig 1D). Participants (N=32) were not in control of the direction they moved during reach training trials. The handle at the end of the robot arm they were grasping was constrained to a force channel, so participants only chose the distance they moved, not direction, removing any performance error. If they attempted to move outside of the pathway, a resistant force, proportional to the depth of penetration with a stiffness of 2 N/mm and a viscous damping of 5 N/(mm/s), was created perpendicular to the pathway (Henriques & Soechting, 2003). During the error clamp phase of the experiment, participants were instructed to actively move their hands thus these trials were identical to the previous two paradigms. Participants still heard the auditory feedback to encourage consistent speed across training paradigms.

#### Localization test trials

All three groups completed a passive localization of their hand position after every training trial. These proprioceptive localization trials (Fig 1E) were executed to one of two targets, 5° on either side of the previous training target. The localization targets were close to the preceding training targets to maximize generalization, but not on the same location to be able to detect if participants simply touched the remembered visual target from the previous trial. All eight hand-targets (55°, 65°, 75°, 85°, 95°, 105°, 115° and 125°; one on each side of each of the training targets) were cycled through before being repeated. After the white arc appeared on the screen, participants’ right unseen, adapted hand was dragged to one of the target locations. Then once their target hand was locked in place, participants used their visible, left index finger, to indicate on the touchscreen, along a 180° arc, where they believed their right, stationary, unseen hand was. The arc was continuously visible until the touchscreen registered the participants estimate. We tested if localization responses were biased towards the preceding visual target in both the end of the aligned and the end of the rotated phase in all three conditions, but there was no bias in 5 of 6 tests. There is a 2.4° bias in the aligned phase of the terminal condition, which is much smaller than the 10° distance between the localization target pairs.

### Data Analysis

We analyzed reach training and hand localization trials separately from each other, but their rates of change (see Table 1) can be compared.

**Table 1.** Adaptation estimates for reach training trials, estimates hand location and a two-rate models slow process prediction. Rate of change estimates, asymptote and average trial participants reached asymptote are provided for each training condition and the corresponding slow process and estimates of hand location. 95% confidence intervals are included for each estimate. Parameters were estimated using an exponential decay model.

| Rate of change |  |  |  |  |
| --- | --- | --- | --- | --- |
|  |  | Continuous | Terminal | Exposure |
| Reach training | rate of change | 27.0%<br>[20.1% - 32.8%] | 14.2%<br>[10.0% - 20.0%] | - |
|  | asymptote | 28.6°<br>[27.8° - 29.5°] | 27.1°<br>[26.1° - 29.7°] | - |
|  | saturation trial | 12<br>[10 - 16] | 19<br>[13 - 27] | - |
| Slow process | rate of change | 3.5%<br>[3.0% - 4.1%] | 3.3%<br>[3.0% - 3.7%] | - |
|  | asymptote | 25.2°<br>[22.5° - 27.2°] | 25.3°<br>[23.0° - 27.2°] | - |
|  | saturation trial | 65<br>[55 - 75] | 72<br>[66 - 80] | - |
| localization | rate of change | 100%<br>[29.0% - 100%] | 43.5%<br>[7% - 100%] | 69%<br>[47% - 100%] |
|  | asymptote | 6.9°<br>[5.9° - 8.0°] | 6.3°<br>[5.3° - 7.8°] | 5.1°<br>[3.8° - 6.4°] |
|  | saturation trial | 1<br>[1 - 7] | 4<br>[1 - 23] | 2<br>[1 - 3] |

### Reaching with a cursor and clamp trials

To quantify reach performance during training, the angular difference between a straight line from the home position to the target and a straight line from the home position and the point of maximum velocity is computed. This was calculated for all training trials both classic and terminal training but only for the error clamp trials for exposure training.

### Hand Localization

Estimates of hand location were based on the angular endpoint error between the movement endpoint of the right unseen hand and the left hands responses on the touchscreen, relative to the home position.

### Analyses

All data was visually screened for incorrect trials. Subsequently, outliers of more than three standard deviations across participants within each trial were also deleted. All measures were normalized, by subtracting out each subjects’ average performance during the second half of the aligned session (e.g. trials 33-64). To see if there were changes in training and test trials, we conducted ANOVAs consisting of a within-subjects factor of trial set and a between-subjects factor of training paradigm. The trial-set factor consisted of four levels: the first 4 rotated trials (early), the final 4 trials from the first rotation (late), the final 4 trials from the second rotation (reversed) and the last 16 trials, to allow for a less noisy estimate, from the clamp phase (clamped). All analyses ignored target location, but each bin of four trials contains a trial to each of the four training targets. Significant main effects and interactions were followed-up by pairwise comparisons, using a Welch t-test and an alpha of .05, where necessary with an FDR correction applied using the p.adjust function in R (Benjamini & Hochberg, 1995).

### Two-Rate Model

We fitted the two-rate model (Smith, Ghazizadeh, & Shadmehr, 2006) to our data. This two-rate model is composed of a slow process that slowly increases over time until it is the driving force of performance, and a fast process that rises quickly but eventually decays back to zero. The sum of these two processes determines the overt behaviour and can explain the rebound seen in the error-clamp phase. During error-clamps, neither process learns, but the fast process will forget how it adapted to the counter rotation, while the slow process still exhibits part of its adaptation from the long initial training, resulting in a rebound.

This model postulates the reaching behavior exhibited on trial t (X_t1_), is the sum of the output of the slow (X_s,t1_) and fast process (X_f,t1_) on the same trial:

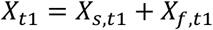

Both processes learn from errors on the previous trial (e_t0_) by means of a learning rate (L_s_ and L_f_), and they each retain some of their previous state (X_s,t0_ and X_f,t0_) by means of their retention rates (R_s_ and R_f_):

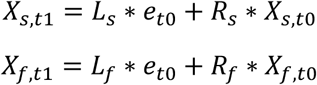

The model is further constrained by making sure the learning rate of the slow process is lower than that of the fast process: L_s_ < L_f_, and by having the retention rate of the slow process be larger than that of the fast process: R_s_ > R_f_. We constrained the parameters to the range [0,1].

All model fitting was done on the mean angular reach deviation at peak velocity during all training reaches, regardless of target angle. The error term was set to zero during the final error clamp phase of the experiment, as the participants did not experience any performance error. The model was fit in R 3.6.1 (R Core Team, 2020) using a least mean-squared error criterion on the six best fits resulting from a grid-search. The parameter values corresponding to the lowest MSE between data and model was picked as the best fit, and this was repeated for continuous and terminal paradigms.

### Rate of Change

We used an exponential decay function with an asymptote to estimate the rate of change for each of the two trial types. The value of each process on the next trial (P_t1_) is the current process’ value (P_t0_) minus the product of the rate of change (L) multiplied by the error on the current trial, which is the difference between the asymptote (A) and the process’ value on the current trial (P_t0_).

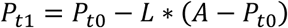

The parameter L was constrained to the range [0,1], and the parameter A to [0,2·max(data)]. For all paradigms using only the first rotations data (trials 65-224), the model was fit to 1) the reach data, 2) the slow process from the two-rate model and 3) localizations. For the latter kind of fit a zero was prepended to account for the fact that responses in these trials already changed through the previous training trial. The parameters were also bootstrapped (1k resamples per fit) across participants to get a 95% confidence interval for both parameters. The first trial where the modelled process based on the group average fell inside the bootstrapped confidence interval for the asymptote is taken as the saturation trial.

The datasets for the current study are available on Open Science Framework, https://osf.io/6q2zd/ while the code and analysis scripts are available on github https://github.com/JennR1990/VisualFeedback.

## Results

We used multiple approaches to investigate if reducing sensory prediction and performance errors during training reduces rate of adaptation in motor learning or slows the rapid changes in estimates of hand location. Specifically, we used a combination of rate of change computations and mixed ANOVAs. Figure 3 shows reach training trials for both the continuous and terminal paradigm, and the error clamp trials for the exposure paradigm. Figure 4 shows all paradigms estimates of hand location during training.

**Figure 3.**
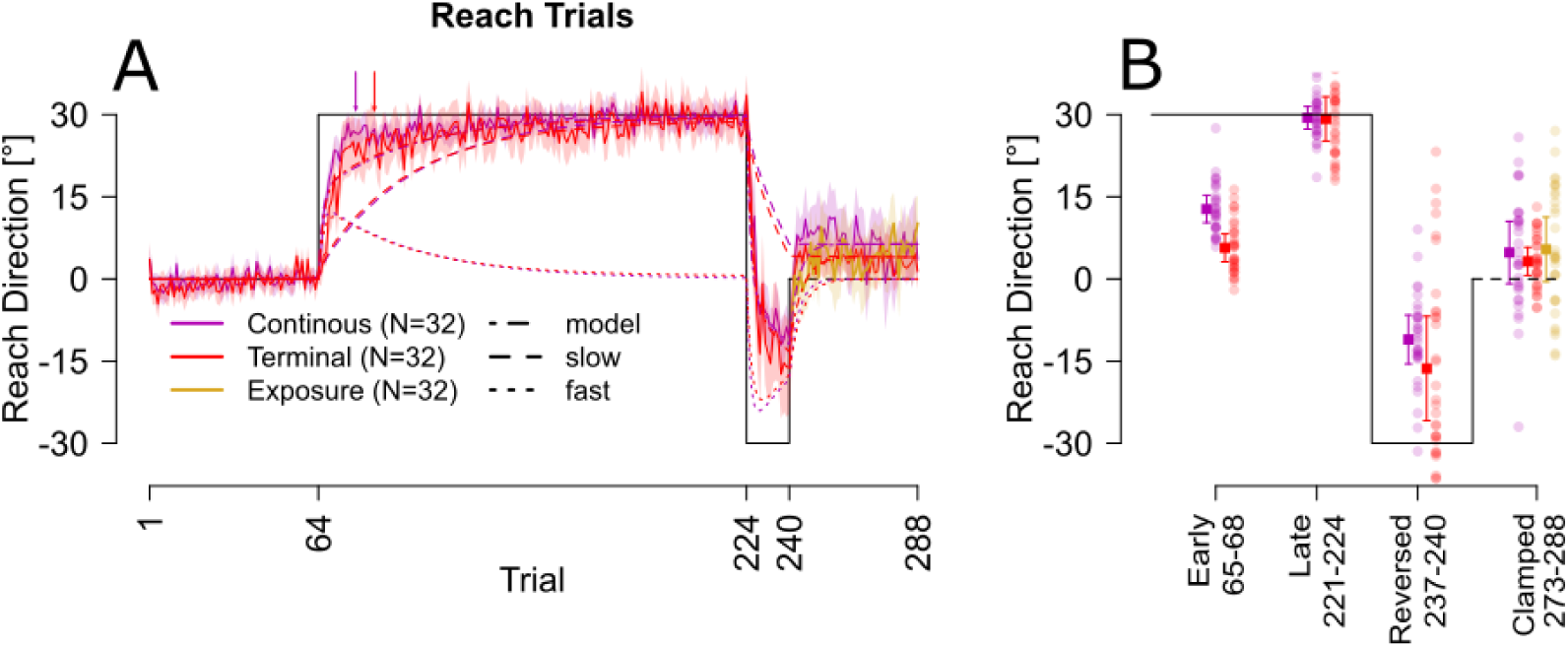
Reach directions during training. **A**. Angle at peak velocity for all three paradigms across the entire training paradigm. Solid lines are the averages across participants within each paradigm, and the corresponding shaded regions are the 95% confidence intervals. Two-rate model estimates are included as dashed and dotted lines in the same color of the paradigms reach data. Colored arrows indicate the average trial at which participants reached asymptote, saturation trial. **B**. Average reach direction during early, late, reversed and EC phases. Individual data is shown around the mean for each paradigm.

**Figure 4.**
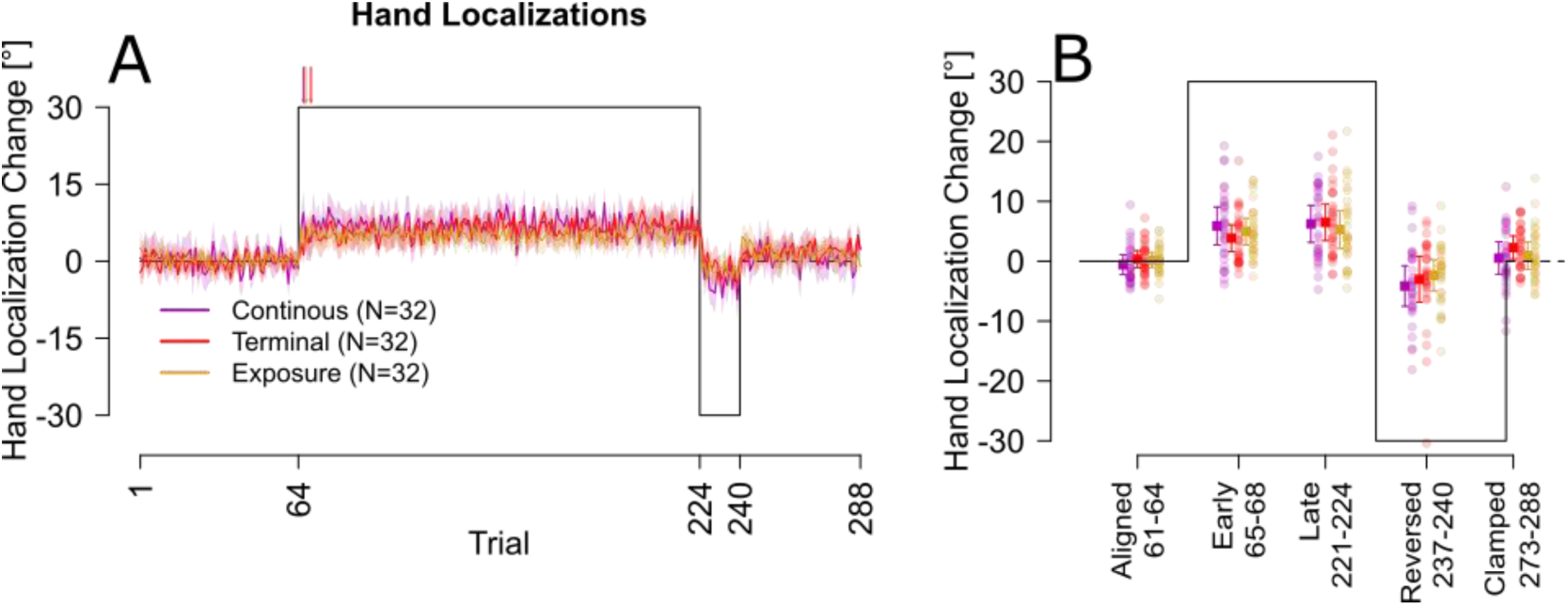
Estimates of Hand location for all training paradigms. **A**. Estimates of hand location throughout the course of training. The solid-colored lines are the deviations between actual and indicated hand position averaged across all participants within a paradigm, the corresponding shaded regions are 95% confidence intervals. Colored arrows indicate the trial participants reached asymptote on average, saturation trial. **B**. Close-up of estimates of hand location for early, late, reversed and error clamped phase. Individual participant data are coded by color along side paradigm averages with error bars representing +/− 2 SE.

### Speed of visuomotor adaptation

Participants in the exposure training paradigm were not in control of movement direction during the first three phases of the experiment and thus were not included in this analysis. Rates of change in the initial learning phase are faster for continuous training [27%, 95%CI: 20.1% - 32.8%] than for the terminal training paradigm [14.2%, 95%CI 10.0% - 20.0%] as shown in Table 1. This is reflected in the average degree of compensation in the early phase of training (Fig 3A) which shows that compensation for terminal feedback (5.68°) was lower than that for continuous feedback (12.77°). This rate of changes meant learning hit an asymptote of 28.6° by the 12^th^ trial for continuous feedback and a similar asymptote of 27.1° by the 19^th^ trial for terminal feedback training (Fig 3A). However, a closer inspection of these trial-by-trial reaches in Fig 3A shows that while continuous feedback reaches maximum compensation by the 12^th^ trial, by that same trial for the terminal feedback shows that compensation is merely a few degrees or 10% behind. This last 10% compensation is what requires the additional 7 training trials to reach a similar asymptote level. Unfortunately, the two-state model can’t capture these small differences, so that the model output, as well as slow and fast process largely overlap as shown in Fig 3A. When we compare the reach deviations for the main blocks of trials (early, late, reversed and clamped) for terminal and continuous training, as plotted in Fig 3B, we find a weak effect of paradigm [F(1,62)=8.04, P=.006, η^2^=.04], which is likely driven by the smaller amount of compensation early in the first rotation [t(59.763)=-1.24, P<.001, η^2^=.36] for the terminal feedback group compared to continuous, as all other block comparisons were non-significant [P>.05]. However, given the large effect of block (due to large changes in cursor rotation), no interaction was found between these two paradigms across these four blocks. Together, these results indicate terminal feedback reduces learning rate, but extent of learning is comparable after 20 trials (see table 1).

All participants, including those in the exposure training group, controlled the movement direction of their hand during the final clamp phase of the experiment. As can be seen in the last block of Fig 3B, we found no difference in the resulting rebound for these final 16 trials of the error clamp phase for the three training paradigms (continuous, terminal and exposure), [F(2,93)=0.47, P=.62]. This indicates that this learning, usually considered a proxy for slow and thus implicit learning, is equally robust across different type of feedback during the training.

### Estimates of hand location

Following every training trial, we measured changes in hand estimate by having participants indicate the felt location of their unseen right hand after it was passively displaced. All training paradigms produced robust shifts in felt hand position ∼6°, (asymptotes in table 1). More importantly, the shift saturated within 1-4 rotated training trials regardless of training. An exponential decay model fit to the localization data indicates that continuous training produces the fastest and largest shift in hand localization. Exposure training and terminal feedback training require a couple of additional trials to reach asymptote. The average shift in felt hand location for exposure training, 5.1° deg, fell below the confidence intervals for the average shifts produces in the other training paradigm (5.9° & 5.3°) as reported in table 1. However, a mixed ANOVA including training paradigm and time point showed no significant effect of training [F(2,93)=0.05, P=0.9] or interaction between training and time point [F(3,279)=1.44, P=.20]. This supports the robustness of shifts in hand localization in response to perturbed visual feedback.

## Discussion

We have previously shown that changes in estimates of unseen hand location shift saturate after a single trial of classical visuomotor adaptation with a continuously visible cursor. Here we measure the extent that this surprisingly rapid saturation may be slowed down with reduced feedback during training, that is, with terminal feedback or with robot constrained movements in an exposure paradigm. By measuring estimates of unseen hand position every other training trial, we captured this implicit component of adaptation in finer detail. We found that even with reduced feedback, changes in felt hand position, saturate very quickly during training, earlier than motor adaptation saturates. Training with terminal feedback or with passive exposure of a 30° rotation only slowed the saturation of these shifts in proprioceptive estimate of hand position by a few trials. Terminal visual feedback produced the slowest rate of change, taking 1-13 trials, compared to 1-7 trials for continuous feedback, but an equally large shift in felt hand position. Exposure training led to an equally rapid rate of change but slightly smaller shifts (5.1° vs 6.3° & 6.9°) than either of the other paradigms. Exposure training does not involve volitional reaches so that a learning rate can not be determined for this condition. However, we identified that reducing feedback to only the endpoint position slowed saturation of motor adaptation to a greater extent, requiring 19 trials for participants to reach saturation for terminal adaptation compared to only 12 trials for classical visuomotor adaptation. Nonetheless, all training paradigms produced equivalent rebounds; as well as comparable two-state parameters. In summary, motor adaptation and changes in felt hand position saturated quickly even when visual and motor signals were reduced during training, with only a small reduction in speed of these changes, demonstrating how rapidly implicit changes can emerge.

### Adaptation to Varying Types of Feedback

Humans are very visually dominant beings and favour vision over many other senses for guiding reaching movements. Thus, it is not surprising that reducing visual feedback of the reach to the end when adapting to a visual perturbation can result in poorer learning performance compared when the cursor is continuously visible. Nonetheless, many studies, including ours, find that given enough training trials similar levels of asymptote are achieved for both training paradigms (Brudner et al., 2016; Heuer & Hegele, 2008; Rand & Rentsch, 2016; Schween & Hegele, 2017; Song, Adams, & Legon, 2020; Wijeyaratnam, Chua, & Cressman, 2019), although in some cases, learning extent is smaller (Barkley et al., 2014) The exact difference in the rate of the learning is not usually measured or reported in previous studies; only a handful of studies compare whether the average first block of trials differ between the different paradigms (e.g. Taylor et al., 2014). In the current study, we fit a single exponential to the two training paradigms, we find that compensation for a terminal feedback visuomotor rotation is only half as fast as that for a continuous distortion (14.2% vs 27%) and takes 30% more training trials (19 vs 12) to saturate. By the 12^th^ trial, however, compensation produced with terminal feedback is only 10% lower than those for continuous. The overall speed and shape of the learning rate could explain conflicting results regarding whether performance in terminal and continuous feedback training paradigms differ. Taken together, this indicates the same mechanisms may facilitate learning in all three conditions, but the reduced feedback merely diminishes the overall speed by which motor and sensory changes hit asymptote levels.

It has been suggested that terminal feedback relies on cognitive strategies early in learning, which is why reaction times are longer and less consistent when reaches with terminal-cursor feedback are compared to continuous feedback (Hinder et al., 2010; Taylor et al., 2014; Wijeyaratnam et al., 2019). However, when we fit a two-rate model, where fast and slow rates have been linked to explicit and implicit components of learning, we find no noticeable differences between the processes for terminal and continuous training and no difference in the size of the rebound for all three paradigms as shown in Fig 3A. If the two-rate model reflects differences in explicit and implicit components, these components do not seem to differ much for the different training paradigms. Moreover, the rebound during the clamped trials emerged and was the same size in the exposure training as for the other training groups. As in our previous studies using exposure training (Cressman & Henriques, 2010; Mostafa et al., 2019; Ruttle et al., 2018; Salomonczyk et al., 2013), this suggests that visual-proprioceptive discrepancies are sufficient to lead to implicit changes in hand movements.

### Learning-induced Changes in Hand Localizations

Following the completion of every training trial participants indicated the felt position of their then passively displaced hand. Shifts in felt hand position have been shown to be implicit (Modchalingam, Vachon, ’t Hart, & Henriques, 2019) and driven by the visual-proprioceptive mismatch between visible cursor location and felt position of the hand (Henriques & Cressman, 2012; Mostafa et al., 2019; Salomonczyk et al., 2013). Previous work in our lab and others has shown that the shift in felt hand position is a robust feature of learning under various conditions (Cameron, Franks, Inglis, & Chua, 2012; Henriques & Cressman, 2012; Izawa & Shadmehr, 2011; Ruttle, ’t Hart, & Henriques, 2021; Ruttle et al., 2018). Here we were able to go a step further by concurrently measuring and modeling these shifts in felt hand position to be able to identify a rate of change and level of asymptote.

As in the continuous-cursor training, the changes in unseen hand location estimates were rapid; with most participants for all groups saturating within a few trials. Nonetheless, terminal feedback required a few additional trials for changes in hand localization to reach a similar asymptote compared to continuous training. In our previous study comparing terminal and continuous feedback training, we found that the proprioceptive recalibration (change in hand estimates) required a third block of 99 trials before achieving the same magnitude of proprioceptive recalibration (Barkley et al., 2014).This is most likely because the method for measuring perceived hand location used in the previous study was a two-alternative force choice (2-AFC) method involving 50 trials to get a single estimate. While the 2-AFC method does an equivalent job of measuring the magnitude of proprioceptive calibration as the method used in this and other studies, it requires a far more training to saturate (Clayton, Cressman, & Henriques, 2014; Ruttle, Cressman, ’t Hart, & Henriques, 2016; Zbib, Henriques, & Cressman, 2016). The method used in the current study is able to measure hand localization shifts much faster with the same consistency (Clayton et al., 2014).These differing methods highlight how sensitive proprioception is to previous exposure to misaligned visual feedback, where only short exposures to differing visual environments produce fast changes in proprioceptive mapping.

Exposure training led to a similar rate of change in hand localization as classical visuomotor training, requiring only one more training trial to reach asymptote. A previous study done by our lab (Ruttle et al., 2018), where we measure changes in hand estimate after every 6-12 cursor-rotation training trials, we could not distinguish difference in rates in these changes between exposure and classical 30° visuomotor training, partly due to the coarser time resolution as well as the larger-than-usual changes in perceived hand position for this exposure training. In this previous paper, the average proprioceptive recalibration for exposure training was 10°, which is larger than the 5°-7° shift usually seen in both our exposure (Cressman & Henriques, 2010; Mostafa et al., 2019; Salomonczyk et al., 2013) and classical training paradigms (Barkley et al., 2014; Modchalingam et al., 2019; Ruttle et al., 2016), including those measured in the current study. However, all these shifts in perceived hand location are within a reasonable range and really emphasize the robustness and rapidness of changes in felt hand position that co-occur when experiencing altered visual feedback of the hand.

## Conclusion

Here we show that implicit changes in felt hand position appear incredibly quick, regardless of available feedback during training with a rotated cursor. Reducing feedback merely slowed down saturation by one or two trials. The impact was greater for reach adaptation, with rate of adaptation for terminal being substantially slower than continuous. We find a similar size in rebound, indicating similar slow learning accrued with continuous, terminal and exposure training. In conclusion, the implicit changes like those estimated with changes in felt hand position, are rapid and resilient feature of adaptation which likely contributes to both early and late learning.

## Notes

https://osf.io/6q2zd/

https://github.com/JennR1990/VisualFeedback.git

### Competing Interest Statement

The authors have declared no competing interest.

